# Nuclear growth and import can be uncoupled

**DOI:** 10.1101/2023.04.19.537556

**Authors:** Pan Chen, Sampada Mishra, Daniel L. Levy

**Affiliations:** Department of Biochemistry and Molecular Biology and Zhejiang Key Laboratory of Pathophysiology, School of Basic Medical Sciences, School of Medicine, Ningbo University, Ningbo, Zhejiang 315211, China; Department of Molecular Biology, University of Wyoming, Laramie, WY 82071, USA

## Abstract

What drives nuclear growth? Studying nuclei assembled in *Xenopus* egg extract and focusing on importin α/β–mediated nuclear import, we show that, while nuclear growth depends on nuclear import, nuclear growth and import can be uncoupled. Nuclei containing fragmented DNA grew slowly despite exhibiting normal import rates, suggesting nuclear import itself is insufficient to drive nuclear growth. Nuclei containing more DNA grew larger but imported more slowly. Altering chromatin modifications caused nuclei to grow less while still importing to the same extent or to grow larger without increasing nuclear import. Increasing heterochromatin in vivo in sea urchin embryos increased nuclear growth but not import. These data suggest that nuclear import is not the primary driving force for nuclear growth. Instead, live imaging showed that nuclear growth preferentially occurred at sites of high chromatin density and lamin addition, whereas small nuclei lacking DNA exhibited less lamin incorporation. Our hypothesized model is that lamin incorporation and nuclear growth are driven by chromatin mechanical properties, which depend on and can be tuned by nuclear import.

## INTRODUCTION

During early embryonic development, nuclear size decreases concomitantly with cell size (Wesley *et al*., 2020). Moreover, abnormal nuclear size and shape are a readout for many diseases, especially the enlarged nuclei present in most cancer cells (Zink *et al*., 2004; Dey, 2010). Thus, how nuclear size is regulated to ensure proper cell function is a fundamental question in cell biology. One notable mechanism responsible for regulating nuclear size involves nuclear import (Levy and Heald, 2010; Kume *et al*., 2017; Brownlee and Heald, 2019; Cantwell and Nurse, 2019; Jevtic *et al*., 2019; Wing *et al*., 2022). The application of novel techniques such as microfluidics, high-throughput screens, and super-resolution microscopy has identified regulators of nuclear envelope (NE) expansion, including importins, the endoplasmic reticulum, nuclear lamins, nucleoplasmin, and phospholipid synthesis (Levy and Heald, 2010; Versaevel *et al*., 2014; Jevtic *et al*., 2015; Edens *et al*., 2017; Kume *et al*., 2017; Brownlee and Heald, 2019; Chen *et al*., 2019; Jevtic *et al*., 2019; Mukherjee *et al*., 2020; Tamashunas *et al*., 2020; Deolal *et al*., 2021; Atanasova *et al*., 2022). Many of these regulators act through nuclear import and are likely conserved, given that these studies were conducted in organisms ranging from fission yeast and sea urchins to *Xenopus* and mammalian cells. Whereas it is clear that modulating nuclear import can tune nuclear size, it is unknown if nuclear import itself is sufficient to drive nuclear growth or if other forces promote expansion of the NE.

An emerging view is that the mechanical properties of chromatin may play a role in nuclear size regulation (Dahl *et al*., 2008; Stephens *et al*., 2019b). Changes in chromatin modifications and organization are associated with altered nuclear morphology (Stephens *et al*., 2018; Stephens *et al*., 2019a; Stephens *et al*., 2019b; Heijo *et al*., 2020; Flores *et al*., 2021). Increasing the amount of heterochromatin using histone demethylase inhibitors concomitantly increased chromatin stiffness and rescued abnormal nuclear shape (Stephens *et al*., 2018). Increasing the DNA content within the nucleus can accelerate nuclear expansion, possibly due to a general increase in overall chromatin stiffness (Heijo *et al*., 2020). We previously identified the histone chaperone nucleoplasmin (Npm2) as a regulator of nuclear size during *Xenopus laevis* embryogenesis and proposed that Npm2 promotes nuclear expansion by increasing chromatin compaction and stiffness through histone import and incorporation into chromatin (Chen *et al*., 2019). Because chromatin has been proposed to behave like a spring, one possibility is that chromatin can transmit mechanical forces to the NE and that the extent of these forces depends on the physical properties of the chromatin (Chalut *et al*., 2012).

One structure that is critical to the regulation of nuclear morphology is the nuclear lamina, a stiff layer of intermediate lamin filaments that lines the inner surface of the NE (Cho *et al*., 2019). For example, nuclei in lamin B1–deficient mice are misshapen, softer, and exhibit nuclear blebs (Coffinier *et al*., 2011). Additionally, the nuclear lamina provides an interaction platform for transcriptionally repressed chromatin, and tethering of chromatin to the nuclear periphery can impart stiffness to the nucleus (Schreiner *et al*., 2015). Whereas chromatin stiffness determines the nuclear mechanical response to small applied forces, the nuclear response to large applied forces is dominated by the mechanical properties of the nuclear lamina (Banigan *et al*., 2017; Stephens *et al*., 2017). Of note, lamins must first be imported into the nucleus and incorporated into the lamina underneath the NE to influence nuclear morphology (Jevtic *et al*., 2015). Fundamental questions remain about how lamin incorporation occurs and whether lamin distribution within the NE affects nuclear size.

Nuclear F-actin is another structure that can influence nuclear architecture (Zahler, 2020). In *Xenopus* oocytes, high levels of nuclear-imported actin stabilize the large nucleus through the formation of a filamentous actin network (Bohnsack *et al*., 2006). In *Xenopus* extract, cultured mammalian cells, and mouse embryos, nuclear F-actin contributes to nuclear growth and the regulation of nuclear morphology (Baarlink *et al*., 2017; Oda *et al*., 2017; Heijo *et al*., 2020; Mishra and Levy, 2022). In starfish oocytes, a transient increase in nuclear F-actin exerts intranuclear forces that drive NE breakdown (Wesolowska *et al*., 2020).

In this study, we sought to understand the interplay between nuclear import and intranuclear structures in driving nuclear growth. *Xenopus* egg extracts are an ideal system with which to address these issues, as these extracts support de novo assembly and growth of nuclei; their biochemical manipulation is straightforward; and they are transcriptionally inert, eliminating confounding effects of altered chromatin structure on transcription (Wang and Shechter, 2016; Wang *et al*., 2019). Cycloheximide is added to arrest the extract in interphase, so there is also no new protein synthesis. Furthermore, inhibiting the nuclear export factor CRM1 in early-stage *Xenopus* embryos did not affect nuclear size (Edens and Levy, 2014) and nuclear export is negligible in the egg and early embryo (Callanan *et al*., 2000), so we can use *Xenopus* egg extract to focus on the contribution of nuclear import without potential complications from nuclear export. Through a variety of different approaches, we have manipulated the amount and structure of the chromatin within the nucleus. Although these manipulations had consistent effects on nuclear size, in many instances there was no correlation between the rates of importin α/β–mediated nuclear import and nuclear growth, suggesting that nuclear import itself is not sufficient to drive nuclear growth. Instead, we hypothesize that chromatin-mediated forces at the NE drive nuclear growth by promoting lamin incorporation and present data that are consistent with this model. Thus, we propose that nuclear import tunes nuclear size through the import of key structural components but is not the primary driving force for nuclear expansion.

## RESULTS

### Nuclear size is sensitive to the amount of DNA

Previous studies have suggested that blocking nuclear import prevents nuclear growth (Levy and Heald, 2010; Chen *et al*., 2019; Jevtic *et al*., 2019). To verify this result, we assembled nuclei in *X. laevis* egg extract and then blocked nuclear import through addition of either wheat germ agglutinin (WGA) to block the nuclear pore complex (NPC) or the importin β binding domain of importin α (IBB), a dominant-negative inhibitor of importin α/β–mediated nuclear import. To quantify nuclear import, the extract was supplemented with a reporter consisting of green fluorescent protein fused to the SV40 nuclear localization signal (GFP-NLS), and live time-lapse imaging was performed. The GFP-NLS reporter includes a glutathione S-transferase (GST) tag and is 56 kDa, which is larger than the selective permeability barrier of the NPC. The GFP-NLS nuclear import rate was quantified as described (Mukherjee *et al*., 2020) (also see Methods). Both WGA and IBB treatments nearly completely inhibited GFP-NLS nuclear import as well as nuclear growth (Fig. S1A-C), confirming that importin α/β–mediated nuclear import is necessary for nuclear growth. Because nuclei failed to expand in the presence of IBB, this indicates that nuclear growth is primarily dependent on importin α/β–mediated nuclear import rather than import of cargos regulated by different importins. For this reason, all of our subsequent measurements of nuclear import rates make use of the GFP-NLS reporter because the SV40 NLS is imported via importin α/β (Lu *et al*., 2021). When we refer to “nuclear import,” we specifically mean importin α/β– mediated import with GFP-NLS import rates serving as a proxy for import of cargos regulated by importin α/β.

We previously showed that chromatin occupies proportionately less of the nuclear space as the nucleus grows and that increasing the compaction and stiffness of the chromatin can promote further nuclear growth (Chen *et al*., 2019). To test if reducing chromatin compaction would have the opposite effect, we supplemented nuclei assembled in *Xenopus* extract with the histone methyltransferase Set9 (Batista and Helguero, 2018) or the DNA methyltransferase inhibitor Zebularine (Yoo *et al*., 2004). As expected, both manipulations reduced chromatin compaction, which was demonstrated experimentally by a proportional increase in the nuclear space occupied by chromatin (Fig. S2A-D and S2F-H). Under this condition in which chromatin was less compact, and presumably softer, the nuclei grew less and reached a smaller size (Fig. S2E and S2I). These data further support the idea that chromatin structure can affect nuclear size.

To more directly assess the effect of chromatin on nuclear size, we treated nuclei assembled in extract with Benzonase to completely degrade the DNA. At different time points after Benzonase addition, nuclei were fixed, visualized by immunofluorescence with an antibody against the NPC, and quantified for NE cross-sectional (CS) area. Previous studies have shown that CS nuclear area is an accurate metric for nuclear size and correlates well with total nuclear surface area and volume (Levy and Heald, 2010; Edens and Levy, 2014; Jevtic and Levy, 2015; Vukovic *et al*., 2016). Benzonase efficiently degraded DNA in these nuclei, and nuclear growth almost completely ceased upon Benzonase addition (Fig. 1A). To test if the initial nuclear size mattered, we allowed nuclei to reach different sizes by incubating nuclei in extract for different lengths of time and then adding Benzonase. Again, compared with controls, Benzonase-treated nuclei were significantly smaller (Fig. 1B). We also performed live time-lapse imaging of nuclei visualized with the imported nuclear marker GFP-NLS. Nuclear growth ceased soon after Benzonase addition (Fig. 1C, Movie 1), corroborating our results with fixed nuclei. In addition, we found that Benzonase-treated nuclei were still intact, as they excluded 150-kDa FITC-dextran (Fig. 1D). Of note, Benzonase-treated nuclei were able to import Lamin B3, the primary lamin isoform present in *Xenopus* eggs, but the lamins were largely nucleoplasmic rather than NE-associated as observed in control nuclei (Fig. 1E). This suggests that DNA is required to promote lamin incorporation into the nuclear lamina and nuclear growth, perhaps through chromatin-mediated forces exerted on the NE.

**Figure 1.**
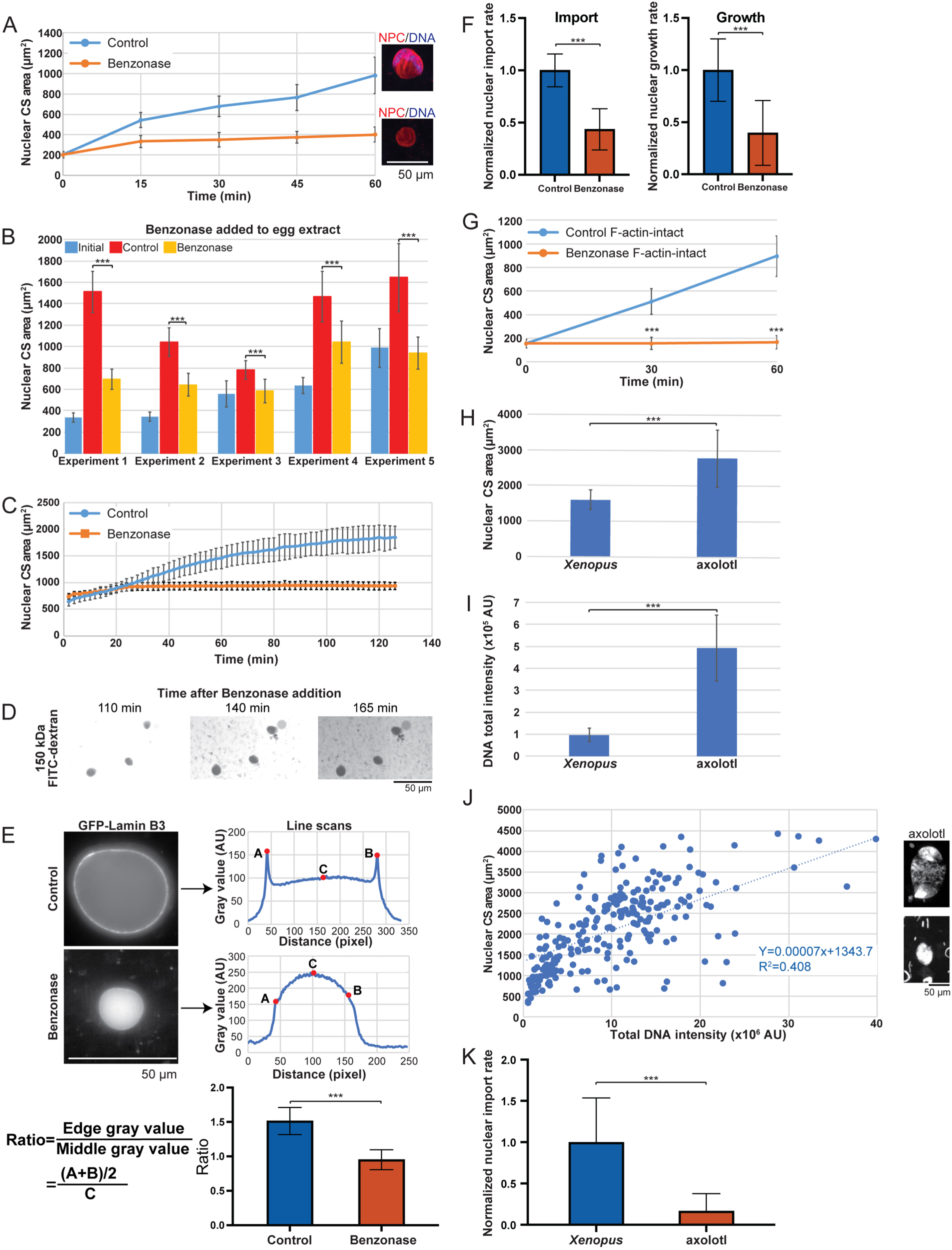
DNA amount influences nuclear growth in *X. laevis* egg extract. **(A-F)** Nuclei were assembled in *X. laevis* egg extract using *X. laevis* sperm chromatin. Assembled nuclei were treated with 2.5 U/µL Benzonase or an equivalent volume of buffer (Control). **(A)** Nuclei were incubated for the indicated lengths of time and visualized by immunofluorescence using mAb414 to label the NPC (red) and Hoechst to label DNA (blue). NPC staining was used to quantify nuclear cross-sectional (CS) area for at least 50 nuclei at each time point. Representative 60-minute images are shown. Scale bar, 50 µm. **(B)** Preassembled nuclei were incubated in extract for different lengths of time (30-60 minutes for Experiments 1–5) to allow nuclei to grow to different initial sizes (Initial) prior to treatment with Benzonase or buffer (Control) for 1 hour. Nuclear size was quantified as in (A) for at least 40 nuclei per condition. **(C)** Preassembled nuclei were treated with Benzonase or buffer (Control) for 10 minutes, and then GFP-NLS (2.9 µM) was added. After an additional 5-minute incubation, live imaging was performed at 2-minute intervals with the same exposure time. Nuclear size was quantified based on the GFP-NLS signal. Data are shown for 9 control nuclei and 5 Benzonase-treated nuclei. **(D)** Preassembled nuclei were treated with Benzonase for 10 minutes, and then FITC-dextran (150 kDa, 4 mg/mL) was added. Live imaging was performed with the same exposure time. Representative images are shown. Scale bar, 50 µm. **(E)** Preassembled nuclei were treated with Benzonase or buffer (Control) for 10 minutes, and then GFP-Lamin B3 (80 nM) was added. After a 90-minute incubation, live imaging was performed with the same exposure time. Representative images are shown. Scale bar, 50 µm. Two Lamin B3 intensity line scans were acquired through the middle of each nucleus in ImageJ Fiji by randomly generating lines and using the “Intensity Profile” tool. To calculate the lamina-to-nucleoplasm Lamin B3 localization ratio, the mean edge intensity (average of points A and B) was divided by the middle intensity (point C). Ratio values greater than one correspond to increased incorporation of Lamin B3 into the nuclear lamina. Representative data from one of two independent experiments are presented. **(F)** After a 10-minute control or Benzonase treatment, GFP-NLS (2.9 µM) was added and incubated for 5 minutes. Live images were then acquired at 2-minute intervals with the same exposure time. The initial nuclear import rate and nuclear growth rate were quantified over 10 minutes as described in the Methods, in each case normalizing to controls. At least 13 nuclei were quantified per condition. **(G)** Nuclei were assembled by adding *X. laevis* sperm chromatin to *X. laevis* egg extract lacking Cytochalasin B (i.e., F-actin-intact egg extract). Nuclei were then treated with 2.5 U/µL Benzonase or an equivalent volume of buffer (Control) for the indicated lengths of time and were visualized by immunofluorescence as in (A). NPC staining was used to quantify nuclear size for at least 50 nuclei at each time point. **(H-K)** Nuclei were assembled in *X. laevis* egg extract using *X. laevis* or axolotl sperm chromatin. After incubation for 2 hours, nuclei were fixed and visualized by immunofluorescence as described in (A). **(H)** Nuclear CS area was quantified for at least 100 nuclei per condition. **(I)** Total DNA intensity was calculated based on Hoechst staining for at least 25 nuclei per condition. **(J)** Nuclear CS area is plotted as a function of total DNA intensity for nuclei assembled with axolotl sperm. Representative images are shown to the right. Scale bar, 50 µm. **(K)** GFP-NLS (2.9 µM) was added to preassembled nuclei and incubated for 5 minutes. Live images were then acquired at 3-minute intervals with the same exposure time. The initial nuclear import rate was quantified over 15 minutes as described in the Methods and normalized to controls. At least 12 nuclei were quantified per condition. Two-tailed Student’s t-tests assuming equal variances: ***p < 0.001. Error bars represent SD.

Nuclear F-actin is known to influence nuclear expansion (Baarlink *et al*., 2017; Mishra and Levy, 2022). Typical *Xenopus* extracts include cytochalasin to depolymerize F-actin, so that could explain why Benzonase-treated nuclei failed to grow. To directly test this idea, we assembled nuclei in actin-intact extracts lacking cytochalasin (Field *et al*., 2017). However, preassembled nuclei lacking DNA still failed to grow in the presence of dynamic F-actin (Fig. 1G). Thus, DNA must be present for nuclear growth to occur.

We next examined whether nuclear size could be altered by an increase in DNA content. In typical extract experiments, nuclei are assembled around *X. laevis* sperm chromatin with a genome size of ∼1.5 Gbp. To increase the DNA content by around 10-fold, we assembled nuclei in *X. laevis* extract using axolotl sperm chromatin with a genome size of ∼16 Gbp. The resulting axolotl sperm–generated nuclei (hereafter, axolotl nuclei) exhibited a larger average CS area by ∼2-fold as compared with nuclei formed around *Xenopus* chromatin (Fig. 1H). Hoechst staining intensity confirmed that the axolotl nuclei contained more DNA than the *Xenopus* nuclei (Fig. 1I). Interestingly, the nucleus-to-nucleus variation in Hoechst staining intensity varied over 10-fold across all axolotl nuclei, perhaps indicating shearing and/or dispersion of axolotl chromosomes prior to nuclear assembly. Fortuitously this provided a wide range of DNA contents to examine. Plotting of nuclear CS area versus DNA intensity for axolotl nuclei revealed a roughly linear correlation (Fig. 1J). Taken together, these data show that nuclear size is sensitive to DNA amount, consistent with previous studies in *Xenopus* (Levy and Heald, 2010; Heijo *et al*., 2020).

### DNA fragmentation leads to smaller nuclear size without affecting nuclear import

During our experiments with Benzonase-treated nuclei, we noted that, in addition to reducing the rate of nuclear growth, benzonase also significantly reduced the GFP-NLS nuclear import rate (Fig. 1F). For this reason, it was unclear if Benzonase reduced nuclear growth due to degradation of DNA and/or reduced nuclear import. We therefore sought conditions that would disrupt DNA structure without impacting nuclear import. We decided to test micrococcal nuclease (MNase) and restriction enzymes, reasoning that RCC1 might be retained in the nucleus under these conditions and support normal nuclear import (Cole and Hammell, 1998). Consistent with this prediction, treatment of preassembled nuclei with MNase led to the digestion of DNA into nucleosomal fragments that remained nuclear (Fig. 2A), and the GFP-NLS nuclear import rate was similar in the MNase-treated and control nuclei (Fig. 2B). In spite of exhibiting similar nuclear import kinetics to controls, MNase-treated nuclei grew more slowly (Fig. 2B-C and Movie 2). As with our Benzonase experiments, we also incubated different sized nuclei with MNase followed by fixation and immunofluorescence. As observed by live imaging, MNase-treated nuclei were smaller than control nuclei regardless of their initial size (Fig. 2D).

**Figure 2.**
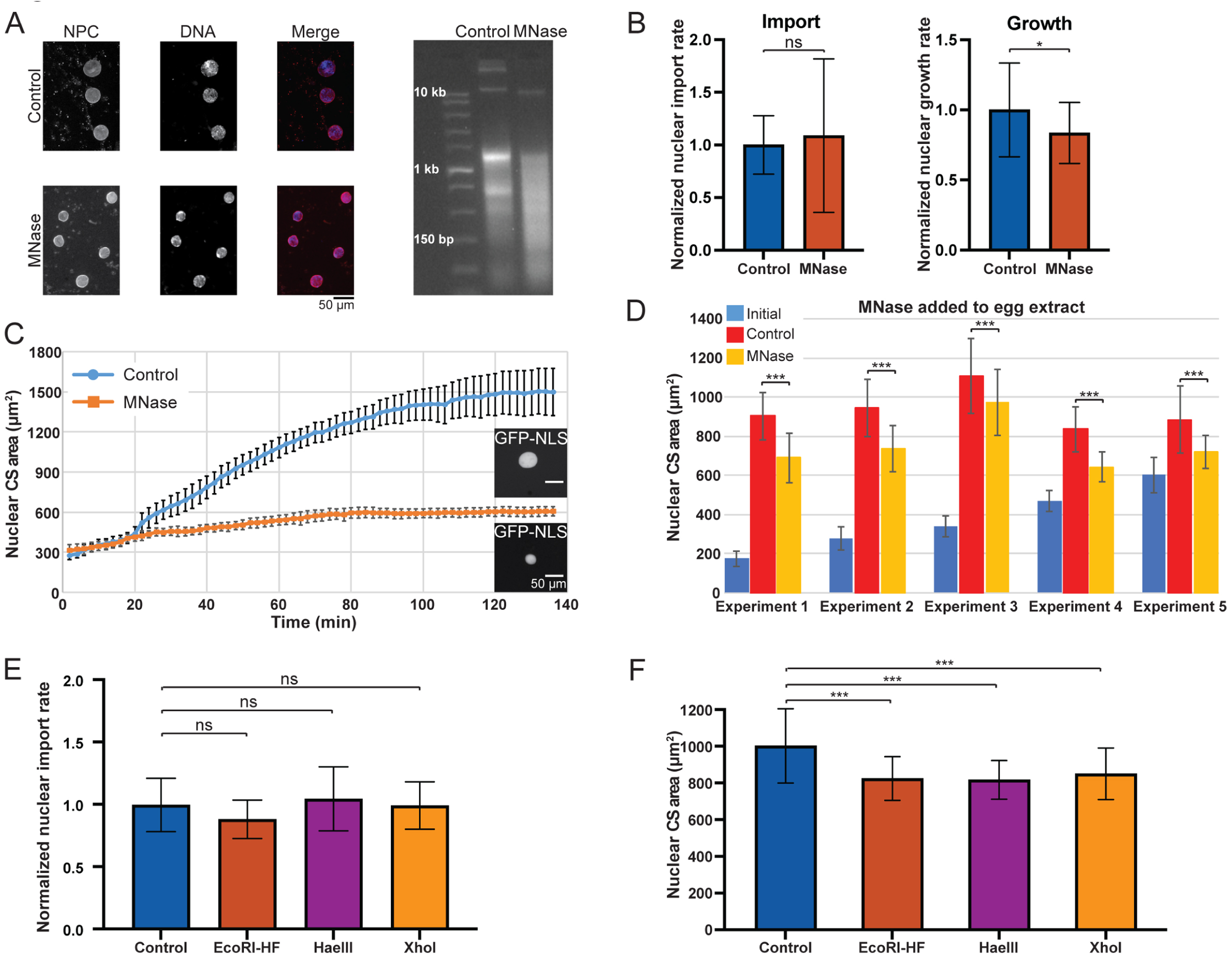
DNA fragmentation reduces nuclear growth but not import in *X. laevis* egg extract. **(A-D)** Nuclei were assembled in *X. laevis* egg extracts with *X. laevis* sperm. Assembled nuclei were treated with 40 U/µL MNase or an equivalent volume of buffer (Control). Similar results were obtained with MNase and MNase-NLS, so those data were combined. **(A)** After a 1-hour incubation with MNase, nuclei were fixed and visualized by immunofluorescence using mAb414 (red) and Hoechst staining (blue). Left: Representative images are shown. Scale bar, 50 µm. Right: DNA was extracted from isolated nuclei, separated on a 2% agarose gel, and stained with ethidium bromide. The left lane shows a DNA ladder with sizes noted. **(B)** After a 10-minute MNase treatment, 2.9 µM GFP-NLS was added and incubated for 5 minutes. Live images were then acquired at 15-sec intervals with the same exposure time. The initial nuclear import rate was quantified over 1.5 minutes as described in the Methods and normalized to controls. Nuclear growth rate was quantified over 10 minutes as described in the Methods and normalized to controls. At least 47 nuclei were quantified per condition. **(C)** After a 10-minute MNase or control treatment, GFP-NLS (2.9 µM) was added and incubated for 5 minutes. Live images were then acquired at 2-minute intervals with the same exposure time. Nuclear CS area was quantified at each time point for 7 control nuclei and 8 MNase-treated nuclei. Representative images are shown at 134 minutes. Scale bars, 50 µm. **(D)** Preassembled nuclei were incubated in extract for five different lengths of time (30-60 minutes for Experiments 1–5) to allow nuclei to grow to different initial sizes (Initial) prior to treatment with MNase or buffer (Control) for 1 hour. Nuclei were then fixed and visualized by immunofluorescence using mAb414 to label the NPC. Nuclear CS area was quantified for at least 40 nuclei per condition. **(E-F)** Nuclei were assembled in *X. laevis* egg extract with *X. laevis* sperm. Assembled nuclei were treated with the indicated restriction enzymes: EcoRI-HF (0.2 U/µL), HaeIII (0.5 U/µL), XhoI (0.2 U/µL), or an equivalent volume of buffer (Control). **(E)** After a 10-minute restriction enzyme treatment, GFP-NLS (2.9 µM) was added and incubated for 5 minutes. Normalized nuclear import rates were quantified as described in (B) for at least 11 nuclei per condition. **(F)** After a 1-hour incubation with restriction enzymes, nuclei were fixed and visualized by immunofluorescence using mAb414 to label the NPC, and the nuclear CS area was quantified for at least 50 nuclei per condition. One set of representative data is shown from three independent experiments. Two-tailed Student’s t-tests assuming equal variances (B and D) and one-way ANOVA tests (E and F): ns, not significant; *p < 0.05; ***p < 0.001. Error bars represent SD.

To further confirm that an intact DNA structure is important for nuclear growth, we used different DNA restriction enzymes to digest the nuclear DNA into smaller fragments, as previously reported in *Xenopus* extract (Kobayashi *et al*., 2002; Heijo *et al*., 2020). We again observed that DNA fragmentation did not affect the GFP-NLS nuclear import rate but that nuclear size was reduced as compared with controls (Fig. 2E-F). These data suggest that chromatin continuity is one aspect of chromatin structure that is important for nuclear growth, independent of nuclear import.

### Chromatin modifications influence nuclear size without affecting nuclear import

Nuclei treated with MNase or restriction enzymes grew less even though nuclear import was unaffected (Fig. 2), suggesting that nuclear import itself is not sufficient to drive nuclear growth. Another instance in which nuclear growth and import were uncoupled was with nuclei assembled around axolotl sperm chromatin, where axolotl nuclei were larger than *Xenopus* nuclei despite exhibiting reduced nuclear import (Fig. 1H and 1K). To further investigate if altering chromatin structure can influence nuclear growth without affecting nuclear import, we manipulated chromatin modifications in *Xenopus* egg extracts using different reagents to alter histone methylation, histone acetylation, or DNA methylation (Miranda *et al*., 2009; Stephens *et al*., 2019b; Esmaeili *et al*., 2020). Potential pleiotropic effects due to altered gene expression could be excluded because *Xenopus* egg extract is transcriptionally inert. Comparing absolute nuclear sizes between treatments was complicated by extract-to-extract variability, so here we present CS nuclear area data normalized to controls. First, we increased the amount of euchromatin using the histone deacetylase inhibitor valproic acid (VPA) or the histone methyltransferase inhibitor 3-Deazaneplanocin A (DZNep) (Stephens *et al*., 2017; Stephens *et al*., 2018; Stephens *et al*., 2019b). Nuclei preassembled in *X. laevis* egg extract were treated with each small molecule for 90 minutes. VPA and DZNep treatment reduced nuclear growth and size relative to controls, leading to 27% and 15% smaller nuclear CS areas, respectively, without affecting the rate of GFP-NLS import (Fig. 3A-D). Thus, even though control and treated nuclei imported to the same extent, treated nuclei grew less than control nuclei, providing further support for the idea that nuclear growth and import can be uncoupled.

**Figure 3.**
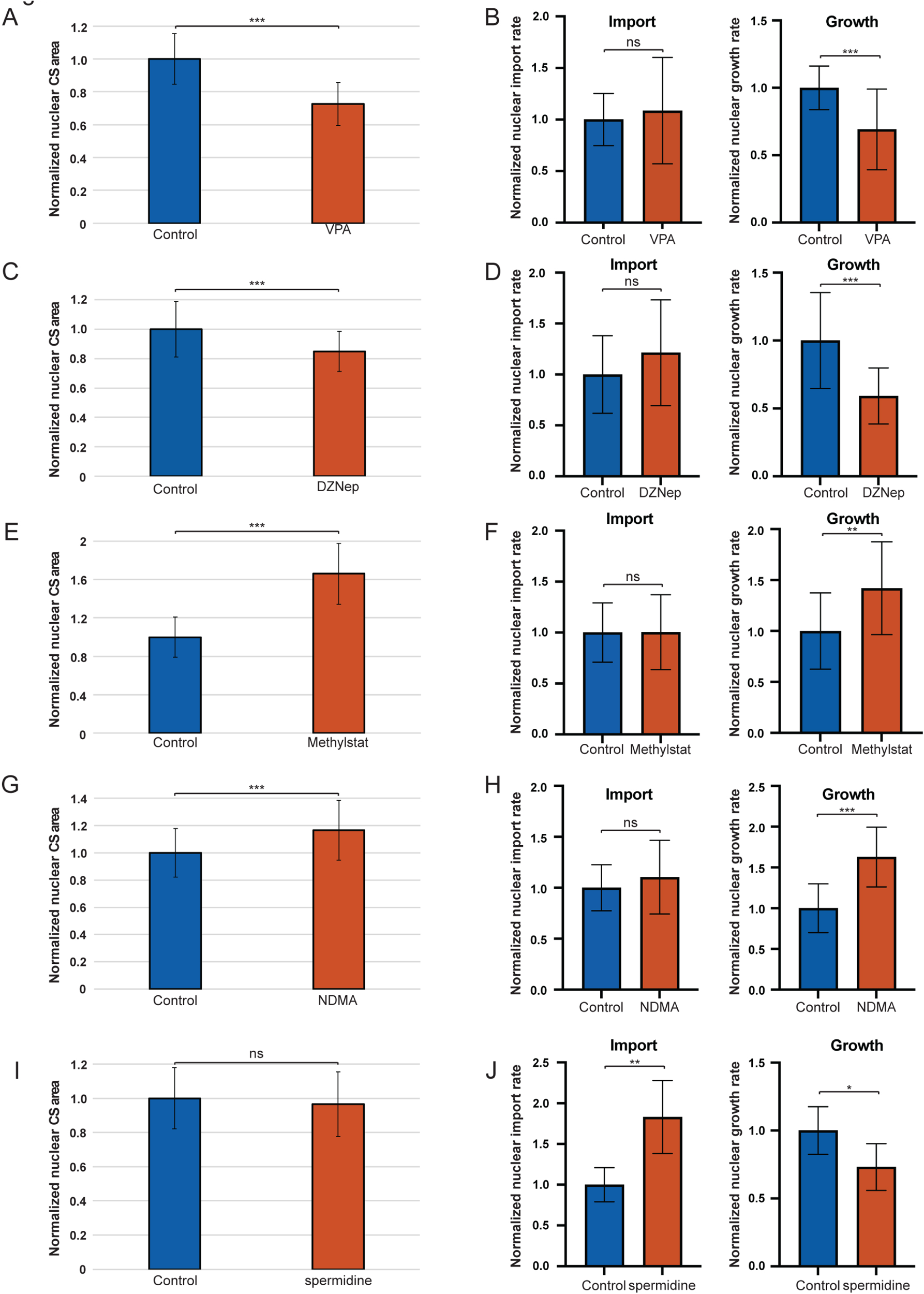
Chromatin modifications can alter nuclear size without affecting nuclear import in *X. laevis* egg extract. Nuclei were assembled in *X. laevis* egg extract with *X. laevis* sperm. **(A-J)** Assembled nuclei were treated as indicated: VPA (0.5 mM) **(A-B)**, DZNep (2 µM) **(C-D)**, Methylstat (4.5 µM) **(E-F)**, NDMA (0.135 M) **(G-H)**, spermidine (10 µM) (**I-J**) or the appropriately matched buffer control (Control). We tested a range of concentrations for each small molecule and here present data for the concentrations that maximally affected nuclear size. **(A,C,E,G,I)** Nuclei were incubated with the indicated small molecules for 90 minutes and were then fixed and visualized by immunofluorescence using mAb414. Nuclear CS area was measured for at least 88 nuclei per condition and normalized to controls. One set of representative data is shown from three independent experiments. **(B,D,F,H,J)** Nuclear import and growth rates for GFP-NLS were quantified as described in Fig. 2B. At least 14 nuclei were quantified per condition except for import measurements with spermidine where 5 nuclei were quantified. Two-tailed Student’s t-tests assuming equal variances: ns, not significant; *p < 0.05; **p < 0.01; ***p < 0.001. Error bars represent SD.

Next, we tested the effect of increasing heterochromatin by supplementing extract with the histone demethylase inhibitor Methylstat (Luo *et al*., 2011; Stephens *et al*., 2018) or the DNA methylator *N*-nitrosodimethylamine (NDMA) (Lin and Hollenberg, 2001; Li and Hecht, 2022). Treatment of nuclei with Methylstat or NDMA led to a 66% or 17% increase in nuclear CS area relative to controls, respectively, and faster rates of nuclear growth without affecting nuclear import rates (Fig. 3E-H). These data show that treated nuclei can grow larger than controls without a concomitant increase in nuclear import. Conversely, low concentrations of spermidine that can subtly affect chromatin conformation (Strickfaden *et al*., 2020) increased nuclear import but reduced the rate of nuclear growth (Fig. 3I-J).

To gain a global picture of how nuclear growth and import correlate, we compiled data for individual nuclei from multiple experiments presented in Figures 2 and 3. While control nuclei exhibited a reasonably strong correlation between nuclear growth and import rates, this correlation was significantly diminished for nuclei in which chromatin structure was altered (Fig. S1D). Taken together, our data suggest that nuclear growth and import can be uncoupled when chromatin structure is modulated. Furthermore, based on what is known about how VPA, DZNep, Methlystat, NDMA, Set9, and Zebularine influence chromatin structure, we propose that an increase in euchromatin leads to smaller nuclei, whereas an increase in heterochromatin promotes nuclear growth (Fig. 3 and Fig. S2), consistent with the notion that stiffer heterochromatin might drive more nuclear growth than softer euchromatin (Chalut *et al*., 2012; Shimamoto *et al*., 2017; Stephens *et al*., 2017; Stephens *et al*., 2018; Chen *et al*., 2019).

### Altering chromatin structure uncouples nuclear growth and import in vivo in sea urchin embryos

To validate our *Xenopus* extract findings in vivo, we performed complementary experiments in *Lytechinus pictus* sea urchin embryos, which are nearly optically clear and allow for live in vivo imaging of nuclear dynamics. A previous study in this system suggested that nuclear growth and import are uncoupled in early stages of development (Mukherjee *et al*., 2020). To further explore this idea, we microinjected one-cell sea urchin embryos with GFP-NLS protein and cultured them in medium supplemented with NDMA. We then quantified nuclear size at the 8- and 16-cell stages. Both the maximum nuclear CS area and initial nuclear growth rate were increased under NDMA treatment as compared with controls (Fig. 4A-C); however, the nuclear import rate was decreased (Fig. 4D). NDMA did not have a large or consistent effect on cell cycle length (Fig. 4E), so the observed effects on nuclear growth and size could not be attributed to extended time spent in interphase. In addition, we confirmed that NDMA altered chromatin structure by showing that mitotic chromosome length was reduced in embryos treated with NDMA (Fig. 4F-G). Thus, these data provide in vivo support that chromatin structure can drive nuclear growth without a concomitant increase in nuclear import, again indicating that nuclear growth and import can be uncoupled.

**Figure 4.**
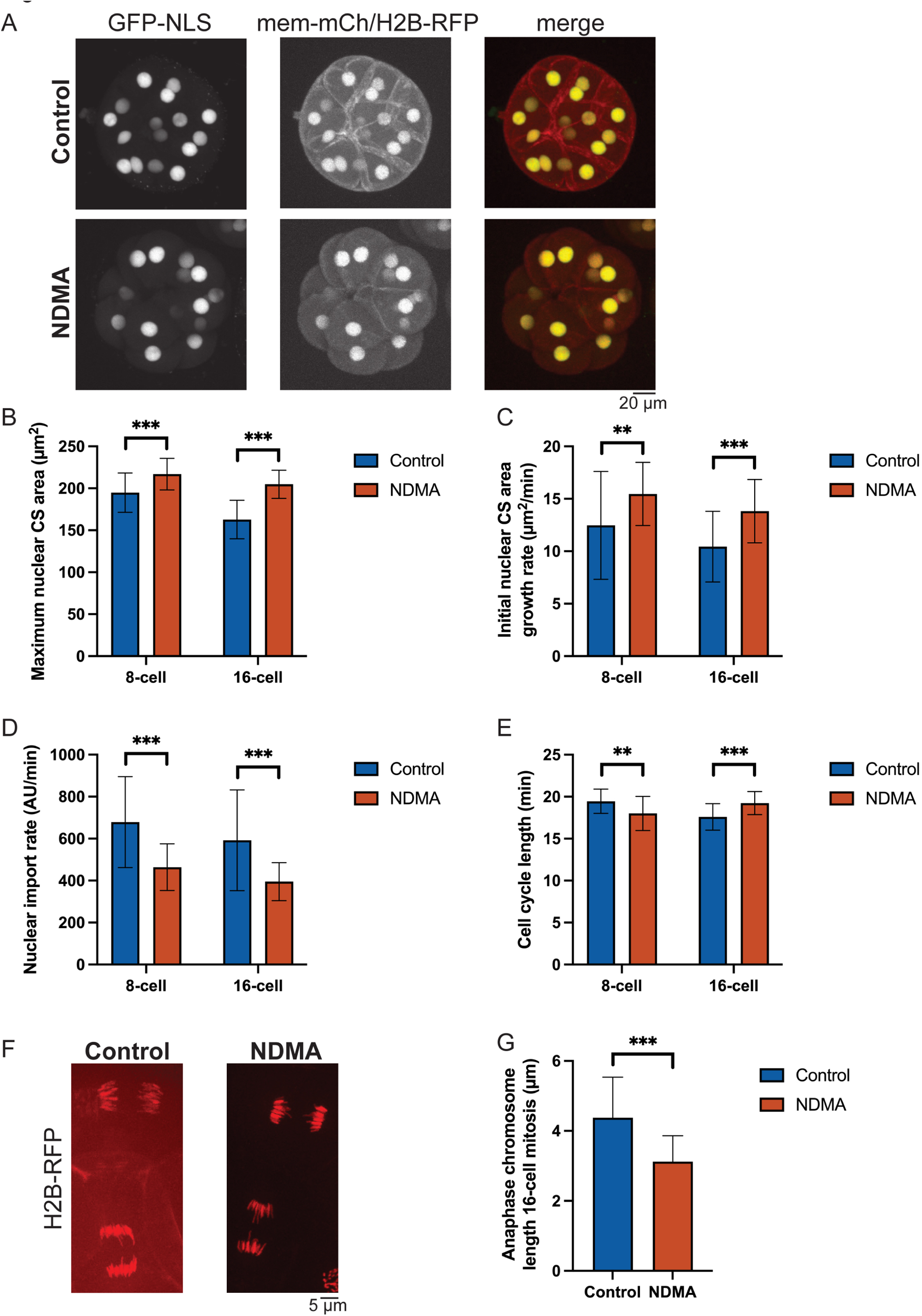
Chromatin modification alters nuclear size in vivo in sea urchin embryos. One-cell sea urchin embryos were microinjected with GFP-NLS protein and mRNAs encoding membrane-mCherry (mem-mCh) to label the plasma membrane and H2B-RFP to label chromatin. Embryos were treated with 64 mM NDMA or an equal volume of buffer. Confocal z-stacks were acquired with a 2-µm step size at 2-minute intervals starting at the 4-cell stage and extending into the 32-cell stage. Quantification focused on the 8- and 16-cell stages, for which the entire cell cycle was imaged. **(A)** Representative maximum-intensity z-projections are shown for 16-cell embryos immediately before entry into mitosis. GFP-NLS is in green. H2B-RFP and membrane-mCherry are in red. Scale bar, 20 µm. **(B)** Nuclear CS areas were measured in the GFP-NLS channel for the entire time-lapses. Because nuclei grow continuously throughout the cell cycle in early embryos, we plotted maximum nuclear CS area within a given embryonic stage to simplify comparisons. **(C)** Initial nuclear CS area growth rates were calculated based on the rate of nuclear growth over 6-10 minutes after the first appearance of nuclear GFP-NLS accumulation. **(D)** Nuclear import rates were calculated based on GFP-NLS import as described in the Methods. **(E)** Cell cycle length, which refers to the time elapsed between consecutive cell divisions, was measured. **(F)** Representative images of anaphase chromosomes from control and NDMA-treated embryos at the 16-cell stage. Scale bar, 5 µm. **(G)** Anaphase chromosome length was measured in embryos at the 16-cell stage, focusing on distinct chromosomes that could be easily visualized. Number of nuclei for (B-E): 51 8-cell control nuclei, 42 8-cell NDMA nuclei, 83 16-cell control nuclei, 53 16-cell NDMA nuclei. Numbers of chromosomes measured for (G): 136 control and 173 NDMA. Cumulative data are shown from 8 control and 7 NDMA-treated embryos. Two-tailed Student’s t-tests assuming equal variances: **p <0.001; ***p < 0.0001. Error bars represent SD.

### Colocalization of nuclear growth, chromatin, and lamin addition

How might chromatin structure influence nuclear growth? It has been suggested that chromatin behaves like a spring, potentially producing intranuclear forces capable of driving nuclear growth (Stephens *et al*., 2019a). We performed time-lapse imaging of chromatin in nuclei assembled in *X. laevis* egg extract. Initially chromatin was dispersed throughout the nucleus but over time tended to aggregate, particularly in proximity to the NE. Concomitantly, chromatin-free regions grew larger in the nuclear interior (Fig. 5A-B and Movies 3-4). This reorganization is reminiscent of reported chromatin phase separation (Fierz and Poirier, 2019; Gibson *et al*., 2019; Bajpai *et al*., 2021). Sites of nuclear growth and blebs often occurred where chromatin was juxtaposed to the NE, as evidenced by neighboring peaks of Lamin B3 and chromatin intensity (Fig 5C), suggesting that accumulation of chromatin at the nuclear periphery may play a role in driving nuclear growth. Consistent with this idea, regions of the NE occupied by the highest density of chromatin tended to represent sites of nuclear growth (Movies 5 and 6) and/or increased NE fluctuations (Movie 7).

**Figure 5.**
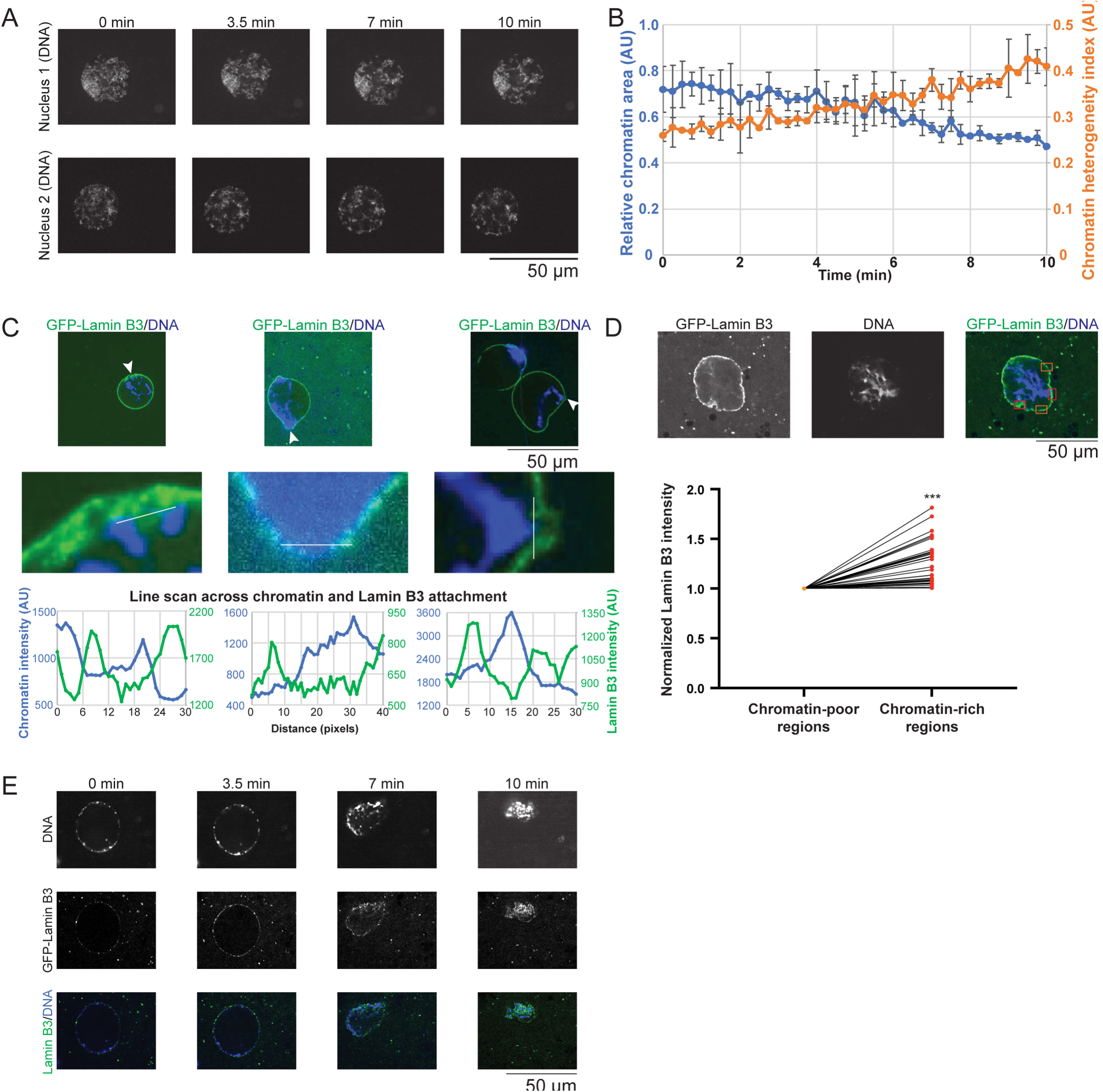
Chromatin colocalizes with sites of nuclear growth and lamin incorporation in *X. laevis* egg extracts. Nuclei were assembled in *X. laevis* egg extract with *X. laevis* sperm. **(A-B)** Preassembled nuclei were labeled with 1 µg/mL Hoechst (DNA). Live time-lapse images were acquired with the same exposure time. **(A)** Time points from two representative time-lapse series are shown. Scale bar, 50 µm. **(B)** Relative chromatin area was calculated as described in Fig. S2A (blue line). To quantify chromatin distribution, we measured the chromatin heterogeneity index as described in (Chen *et al*., 2019). Briefly, for each nucleus two Hoechst intensity line scans were acquired through the middle of the nucleus. The SD of all intensity values along each line was calculated and normalized to the average intensity to obtain the chromatin heterogeneity index (orange line). Larger values correspond to a more heterogeneous chromatin distribution. **(C)** Preassembled nuclei were labeled with 1 µg/mL Hoechst (DNA) and 81.6 nM GFP-Lamin B3. Live images were acquired with the same exposure time. Arrowheads indicate the region shown at 10× magnification in the lower set of images. Intensity line scans were acquired where indicated (blue for chromatin and green for Lamin B3). **(D)** Preassembled nuclei were labeled with 1 µg/mL Hoechst (DNA) and 81.6 nM GFP-Lamin B3. Live images were acquired 10 minutes after Hoechst and GFP-Lamin B3 addition. Representative images are shown. As indicated, multiple rectangular regions of interest were selected randomly for NE regions that were clearly chromatin-rich (red) or chromatin-poor (orange). GFP-Lamin B3 intensity within these rectangles was measured. For each nucleus, GFP-Lamin B3 intensities were normalized to the average intensity within the chromatin-poor regions. Each line represents an individual nucleus. Data are shown for 25 nuclei based on three experiments. **(E)** Preassembled nuclei were labeled with 1 µg/mL Hoechst (DNA) and 44.5 nM GFP-Lamin B3. Live time-lapse images were acquired with the same exposure time. Time points from a representative time-lapse series are shown. Scale bar, 50 µm. Nonparametric Wilcoxon Signed Rank Test: ***p < 0.001. Error bars represent SD.

Based on our observation that small DNA-free nuclei exhibited reduced incorporation of Lamin B3 into the NE (Fig. 1E), we hypothesize that chromatin-mediated pushing forces against the inner NE might facilitate lamin incorporation into the nuclear lamina, thereby driving nuclear growth. To examine this idea, we supplemented extract with GFP-Lamin B3 and performed time-lapse imaging. Regions of the NE with high densities of chromatin incorporated on average 25% more Lamin B3 than chromatin-poor regions of the NE (Fig. 5C-D and Movies 6 and 8), and the lamina often appeared to be more dynamic (Movies 5-7). These data suggest that nuclear growth preferentially occurs at sites of high chromatin density and lamin addition. Consistent with this coupling of the chromatin to the nuclear lamina, extract nuclei infrequently underwent catastrophic rupture in which the chromatin and lamina concomitantly collapsed (Fig. 5E and Movie 9). Based on these observations, we hypothesize a model in which peripheral chromatin both supports the NE and promotes lamin incorporation thereby leading to nuclear growth, although validation of this model will require further experimentation.

## DISCUSSION

In this study we address a fundamental cell biological question about the mechanism of nuclear expansion. Our data support the hypothesis that chromatin amount, continuity, and modifications can influence nuclear growth without necessarily affecting the rate of importin α/μ–mediated nuclear import. Previously, it was reported that an increase in the amount of DNA does not significantly increase nuclear size in *Xenopus* and fission yeast (Neumann and Nurse, 2007; Levy and Heald, 2010). One possible explanation is that the chromatin structure was not substantially changed in those experiments due to either a modest increase in the amount of DNA and/or an imbalance between the amount of DNA and chromatin-binding factors that influence chromatin mechanics.

By altering chromatin organization, we were able to show that nuclear growth and import can be uncoupled, although importin α/μ–mediated nuclear import is required for nuclear growth. We hypothesize that nuclear import provides the intranuclear proteins necessary to drive nuclear growth. For example, nucleoplasmin-mediated import of histones helps to generate a chromatin structure capable of promoting NE expansion (Chen *et al*., 2019) and importin-mediated import of lamins provides the NE building blocks required to expand and stabilize the nuclear membrane (Newport *et al*., 1990; Jenkins *et al*., 1993; Levy and Heald, 2010; Jevtic *et al*., 2015). This model is consistent with our previous study showing that simply increasing the intranuclear protein concentration is insufficient to increase nuclear size (Chen *et al*., 2019). We propose that nuclear import itself is not the primary driving force for nuclear growth, but rather it is the import of specific proteins that in turn generate intranuclear structures capable of producing forces that drive nuclear expansion. Nonetheless, physiological changes in nuclear import can tune nuclear size, for instance through the regulation of nuclear transport factors and NPC numbers (Wesley *et al*., 2020; Chen and Levy, 2023).

We hypothesize that chromatin mechanics are a critical determinant of nuclear expansion and that intranuclear chromatin-mediated forces pushing against the NE promote lamin incorporation and stable nuclear growth. This model can explain why nuclear growth is reduced when DNA is fragmented, a manipulation that reduces chromatin rigidity. It can also explain why nuclear growth is sensitive to various histone and DNA modifications that influence chromatin mechanics. While compact chromatin is more rigid, it may seem counterintuitive that increasing chromatin compaction can result in larger nuclei. Other studies have shown that extreme chromatin decompaction can increase nuclear size (Bustin and Misteli, 2016); however, we argue that the changes in chromatin compaction induced in our study are more subtle such that there could be a physiological compaction range that drives nuclear growth while further compaction leads to smaller nuclei (Chen *et al*., 2019). Another related issue is the distribution of chromatin within the nucleus. Compaction of chromatin at the NE periphery, as observed in *Xenopus* nuclei, may generate chromatin with the necessary mechanics and localization to produce forces at the NE that drive nuclear growth (Chen *et al*., 2019; Stephens *et al*., 2019a). In contrast, highly compacted chromatin that more homogeneously occupies the entire nuclear volume may have the opposite effect and produce smaller nuclei due to reduced chromatin mobility and an inability to generate productive outward pushing forces (Stephens *et al*., 2017; Strickfaden *et al*., 2020). Direct evidence for this model will require comparative measurements of the biophysical properties associated with chromatin that is highly compacted genome-wide versus more subtle localized chromatin compaction. Another area for future research will be to investigate the underlying mechanisms that drive chromatin compaction at the nuclear periphery, in particular the potential contribution of phase separation (Palikyras and Papantonis, 2019; Narlikar, 2020).

Chromatin structure and nuclear size are critical for normal cellular function and development (Carone and Lawrence, 2013; Tsang *et al*., 2020; Ahn *et al*., 2021). Our findings about how chromatin structure influences nuclear expansion add to a growing literature on the diverse mechanisms that regulate nuclear size. Normal nuclear morphology is determined by a balance of forces, including extracellular forces, connections between the cytoskeleton and nucleus mediated by LINC complexes, and intranuclear forces produced by chromatin, F-actin, and intermediate filaments such as lamins (Uhler and Shivashankar, 2018). Disruptions in these forces can affect nuclear size. For example, during normal *Caenorhabditis elegans* aging and advanced aging in progeria, loss of peripheral heterochromatin is associated with changes in nuclear shape (Haithcock *et al*., 2005). With respect to disease, nuclear morphometrics and chromatin condensation state are used as cancer biomarkers (Damodaran *et al*., 2019; Vukovic *et al*., 2021). Future challenges include the integration of known mechanisms of nuclear size control into a unified model and the elucidation of the relative contributions of altered chromatin state versus altered nuclear morphology to disease pathology.

## MATERIALS AND METHODS

### *Xenopus* egg extract and nuclear assembly

*X. laevis* metaphase-arrested egg extract was prepared as described (Maresca and Heald, 2006). Briefly, freshly prepared egg extract was supplemented with LPC (10 µg/mL each leupeptin, pepstatin, and chymostatin), 0.1 mg/mL cytochalasin D, and energy mix (3.8 mM creatine phosphate disodium, 0.5 mM ATP disodium salt, 0.5 mM MgCl_2_). Demembranated *X. laevis* and axolotl sperm chromatin were prepared as described (Murray, 1991; Hazel and Gatlin, 2018). De novo nuclear assembly was performed as described (Chen and Levy, 2018). In brief, interphase-arrested egg extract was generated by supplementation with 0.6 mM CaCl_2_ and 0.15 mg/mL cycloheximide followed by a 25-minute room temperature incubation. A small aliquot of extract was used to test for de novo nuclear assembly quality. All *Xenopus* procedures and studies were conducted in compliance with the US Department of Health and Human Services Guide for the Care and Use of Laboratory Animals. Protocols were approved by the University of Wyoming Institutional Animal Care and Use Committee (Assurance #A-3216-01).

### Recombinant proteins

Recombinant GST-GFP-NLS, GFP-Lamin B3, and IBB were expressed and purified as described (Levy and Heald, 2010; Chen *et al*., 2019). The pK19pA-MN plasmid was a gift from Steven Henikoff (Addgene plasmid #123461; http://n2t.net/addgene:123461; RRID:Addgene_123461). An NLS sequence was introduced into this plasmid before the stop codon using Gibson assembly cloning to generate pK19pA-MN-NLS (pDL125). The inserted sequence was 5′-GGCGGTGGTGGCTCTGGGGGCGGGGGCTCGGGTGGTGGGGGCTCA (linker) CACCATCACCATCACCAT (His-tag) GGCGGTGGTGGCTCTCCACCAAAAAAAAAAAGAAAAGTT (NLS)-3′. Recombinant MNase-NLS protein was expressed and purified as described (Meers *et al*., 2019).

His-tagged Set9 protein expression was induced overnight at room temperature with 1 mM IPTG using BL21(DE3) *Escherichia coli* transformed with pET28 Set9 plasmid (Addgene plasmid #24082). Cells were pelleted at 7000 rpm for 20 minutes at 4°C using rotor JA-10, suspended in lysis buffer (50 mM NaH_2_PO_4_, 300 mM NaCl, 10 mM imidazole, half tablet of Protease Inhibitor Cocktail [Sigma S8830], 0.5 mM PMSF, 30 mg lysozyme, pH 8.0), and incubated on ice for 15 minutes. The cells were sonicated, and cell debris was pelleted at 15000 rpm for 20 minutes at 4°C using rotor JA-20. The supernatant was incubated with 1 mL of Ni-NTA Resin (Thermo Scientific 8822) at 4°C for 2 hours on a rotator. The resin was washed using wash buffer (50 mM NaH_2_PO_4_, 300 mM NaCl, 20 mM imidazole, pH 8.0). Set9 protein bound to the resin was eluted using elution buffer (50 mM NaH_2_PO_4_, 300 mM NaCl, 250 mM imidazole, pH 8.0). Set9 protein was dialyzed into XB buffer (100 mM KCl, 0.1 mM CaCl_2_, 1 mM MgCl_2_, 50 mM sucrose, 10 mM HEPES, 5 mM EGTA) overnight at 4°C. Final protein concentration was measured by Bradford assay.

### Reagents for studying chromatin, DNA, and nuclear import

DZNep (MedChemExpress, HY-12186), Methylstat (Sigma, SML0343-5MG), VPA (MedChemExpress, HY-10585), or NDMA (Sigma, N7756) was added to *X. laevis* egg extract after nuclear assembly. Zebularine (Sigma, Z4775) was added prior to nuclear assembly. For digesting nuclear DNA, 2.5 U/µL Benzonase (Sigma, E1014), 40 U/µL MNase (NEB, M0247S), 0.2 U/μL EcoRI (NEB, R0101S), 0.2 U/μL XhoI (NEB, R0101S), or 0.5 U/μL HaeIII (NEB, R0108S) was added to *X. laevis* egg extract after nuclear assembly. WGA (Sigma, L9640) and 150-kDa FITC-dextran (Sigma, 69658) were used where indicated.

### Immunofluorescence

Immunofluorescence was performed on nuclei isolated from *Xenopus* egg extracts as described (Edens and Levy, 2014, 2016). Briefly, extract containing nuclei was mixed with 20 volumes of fix buffer (50 mM KCl, 2.5 mM MgCl_2_, 250 mM sucrose, 10 mM HEPES [pH 7.8], 15% glycerol, 2.6% paraformaldehyde) and rotated for 15 minutes at room temperature. The solution was layered over 5 mL of cushion buffer (100 mM KCl, 0.1 mM CaCl_2_, 1 mM MgCl_2_, 250 mM sucrose, 10 mM HEPES [pH 7.8], 25% glycerol), and nuclei were spun onto 12-mm circular coverslips at 1000*g* for 15 minutes at 16°C. Nuclei on coverslips were post-fixed in cold methanol for 5 minutes and rehydrated in 0.1% NP-40 in phosphate-buffered saline (PBS). Coverslips were blocked with 3% BSA in PBS overnight at 4°C. They were then incubated at room temperature with primary and secondary antibodies diluted in 3% BSA in PBS for 1 hour each, and DNA was stained with 10 µg/mL Hoechst or EthD-2 for 5 minutes. After each incubation, six washes were performed with 0.1% NP-40 in PBS. Coverslips were mounted on glass slides with Vectashield mounting medium (Vector Laboratories, H-1000) and sealed with nail polish. The main primary antibody used was mouse mAb414 at 1:1000 (BioLegend, #902901), which recognizes NPC FG-repeats. Secondary antibodies included 1:1000 dilutions of Alexa Fluor 488– and 568–conjugated anti–mouse IgG (Molecular Probes, A-11001 and A-11004, respectively).

### Microscopy and image analysis

Wide-field microscopy was performed using an Olympus BX63 upright wide-field epifluorescence microscope equipped to perform multimode, time-lapse imaging using an X-Cite 120LED illumination system. Images were acquired with a high-resolution Hamamatsu ORCA-Flash4.0 digital CMOS camera at room temperature. Confocal microscopy was performed using an Olympus IX83 spinning disk confocal microscope (Yokogawa CSW-W1 SoRA) equipped with an ORCA-Fusion Digital CMOS camera. Olympus objectives included the PLanApoN 2ξ (NA 0.08, air), UPLanFLN 20ξ (NA 0.5, air), UPLanSApo 40ξ (NA 1.25, silicon oil), and UPLanSApo 100ξ (NA 1.35, silicon oil) objectives. Positioning along the x, y, and z axes was controlled by a fully motorized Olympus stage. Acquisition and automation were controlled by Olympus cellSens imaging software, and image analysis was performed using Metamorph software (Molecular Devices).

Images for measuring fluorescence intensity in an individual experiment were acquired using the same exposure time. Total fluorescence intensity and cross-sectional (CS) nuclear area were measured from original thresholded images using Metamorph software. We were confident in our use of CS area to quantify nuclear size, as previous data showed that CS area accurately predicts total nuclear surface area and volume as measured from confocal z-stacks (Levy and Heald, 2010; Edens and Levy, 2014; Jevtic and Levy, 2015; Vukovic *et al*., 2016). For live imaging, nuclei were typically visualized with GFP-NLS or GFP-Lamin B3, whereas fixed nuclei spun onto coverslips were visualized with mAb414. For publication, images were cropped and pseudocolored using ImageJ Fiji (Schindelin *et al*., 2012), but were otherwise unaltered.

Nuclear import was quantified following published protocols (Chen and Levy, 2018; Mukherjee *et al*., 2020). Briefly, *Xenopus* egg extract was supplemented with GFP-NLS protein, and time-lapse imaging was performed. We measured the mean nuclear GFP-NLS pixel intensity for each nucleus and multiplied that value by the nuclear volume to obtain the total intranuclear GFP-NLS signal. These total intranuclear GFP-NLS intensity values were plotted as a function of time, a best-fit linear regression was applied, and the initial slope was used to calculate the initial nuclear import rate. For a given experiment, import rates were normalized to controls to facilitate the compilation of data from multiple independent experiments. To measure nuclear growth rates from these same movies, nuclear CS area was quantified at each time point. Nuclear CS areas were plotted as a function of time, a best-fit linear regression was applied, and the initial slope was used to calculate the nuclear growth rate. For a given experiment, nuclear growth rates were normalized to controls to facilitate the compilation of data from multiple independent experiments.

### Sea urchin experiments

Sea urchin husbandry, embryo generation, and microinjection were as described (Ettensohn *et al*., 2004). Briefly, white painted sea urchins (*Lytechinus pictus*) were obtained from Marinus Scientific and kept at 14–15°C in an aquarium for several weeks in artificial seawater (Reef Crystals; Instant Ocean). Gametes were collected by intracoelomic injection of 0.5 M KCl. Sperm were collected dry and kept at 4°C for up to 1 week. Eggs were rinsed twice, kept at 16°C, and used on the day of collection. The jelly coat of unfertilized eggs was removed by passing them through a 100-μm Nitex mesh (Genesee Scientific) three times to facilitate egg adhesion on protamine-coated glass-bottom dishes (MatTek Corporation). Unfertilized eggs were transferred to protamine-coated glass-bottom dishes for a maximum time of 15 minutes before microinjection and fertilization. Microinjections were performed using a Narishige programmable injector (IM-300). Injection pipettes consisted of borosilicate glass capillaries (1 mm in diameter) pulled with a needle puller (P-1000; Sutter Instruments) and ground with a 30° angle on a needle beveler (BV-10; Sutter) to obtain a 5- to 10-μm aperture. Injection pipettes were back-loaded with 2 μL of appropriately diluted protein and RNA before each experiment and were not reused. Injection volumes were generally <5% of the egg volume, ∼2–5 pL. GST-GFP-NLS protein was diluted to ∼2.5 mg/mL in PBS before injecting. Injecting undiluted GST-GFP-NLS did not affect developmental progression, suggesting that this protein concentration was not detrimental to the embryos (Mukherjee *et al*., 2020). RNA was diluted to ∼50 ng/µL in PBS before injecting. These concentrations were empirically selected to optimize imaging while not affecting developmental progression (Mukherjee *et al*., 2020).

Recombinant GST-GFP-NLS was expressed and purified as described (Levy and Heald, 2010). The membrane-mCherry pCS2+ construct was a gift from Michael Lampson (University of Pennsylvania). The H2B-RFP pCS107 plasmid (pRH199) was a gift from Rebecca Heald (University of California, Berkeley). Each plasmid was linearized with NotI, and mRNA was expressed from the SP6 promoter using the mMessage mMachine kit (Ambion) and isolated in water. RNA stock concentrations were generally ∼700–800 ng/µL. As noted, embryos were treated with 64 mM NDMA (Sigma, N7756) or an equal volume of buffer by direct addition to the culture medium.

Live time-lapse imaging was performed using a spinning disk confocal microscope. Quantification was performed using ImageJ as described (Mukherjee *et al*., 2020). To quantify nuclear size, the maximum CS nuclear area was measured using GFP-NLS, and segmentation was based on gradient detection by thresholding in ImageJ. To quantify nuclear import, we first measured the mean nuclear GFP-NLS pixel intensity and multiplied that value by the nuclear volume to obtain the total intranuclear GFP-NLS signal for each nucleus. These intensity values were plotted as a function of time, and the initial slope was used to calculate the initial nuclear import rate. Import rates were normalized to the cytoplasmic GFP-NLS signal present during the preceding mitosis. Anaphase chromosome length was measured from time-lapse imaging of embryos expressing H2B-RFP. Chromosome length was measured only in cases where distinct chromosomes could be easily visualized.

### Statistical analysis

Averaging and statistical analysis were performed for independently repeated experiments. In general, two-tailed Student’s t-tests assuming equal variances were performed with Minitab 18 to evaluate statistical significance. As appropriate, other statistical tests were applied where indicated using GraphPad Prism. For each immunofluorescence coverslip, at least 100 nuclei were usually quantified. The p-values, number of independent experiments, number of nuclei quantified, and error bars are denoted in the figure legends.

## ACKNOWLEDGMENTS

We thank Dr. Rachel Mueller (Colorado State University) for providing axolotl sperm. We thank Amy Fluet for manuscript editing. This work was supported by the National Institutes of Health/National Institute of General Medical Sciences (R35GM134885 and P20GM103432). PC also acknowledges support from the Zhejiang Provincial Natural Science Foundation of China (no. LQ23C070003).

## AUTHOR CONTRIBUTIONS

Conceptualization: PC, DLL; Investigation: PC (all experiments except for those noted for SM and DLL), SM (Set9 and Zebularine experiments), DLL (sea urchin experiments); Writing – Original Draft: PC; Writing – Review & Editing: PC, SM, DLL; Funding Acquisition: DLL; Supervision: DLL

## SUPPLEMENTAL FIGURE LEGENDS

**Figure S1.**
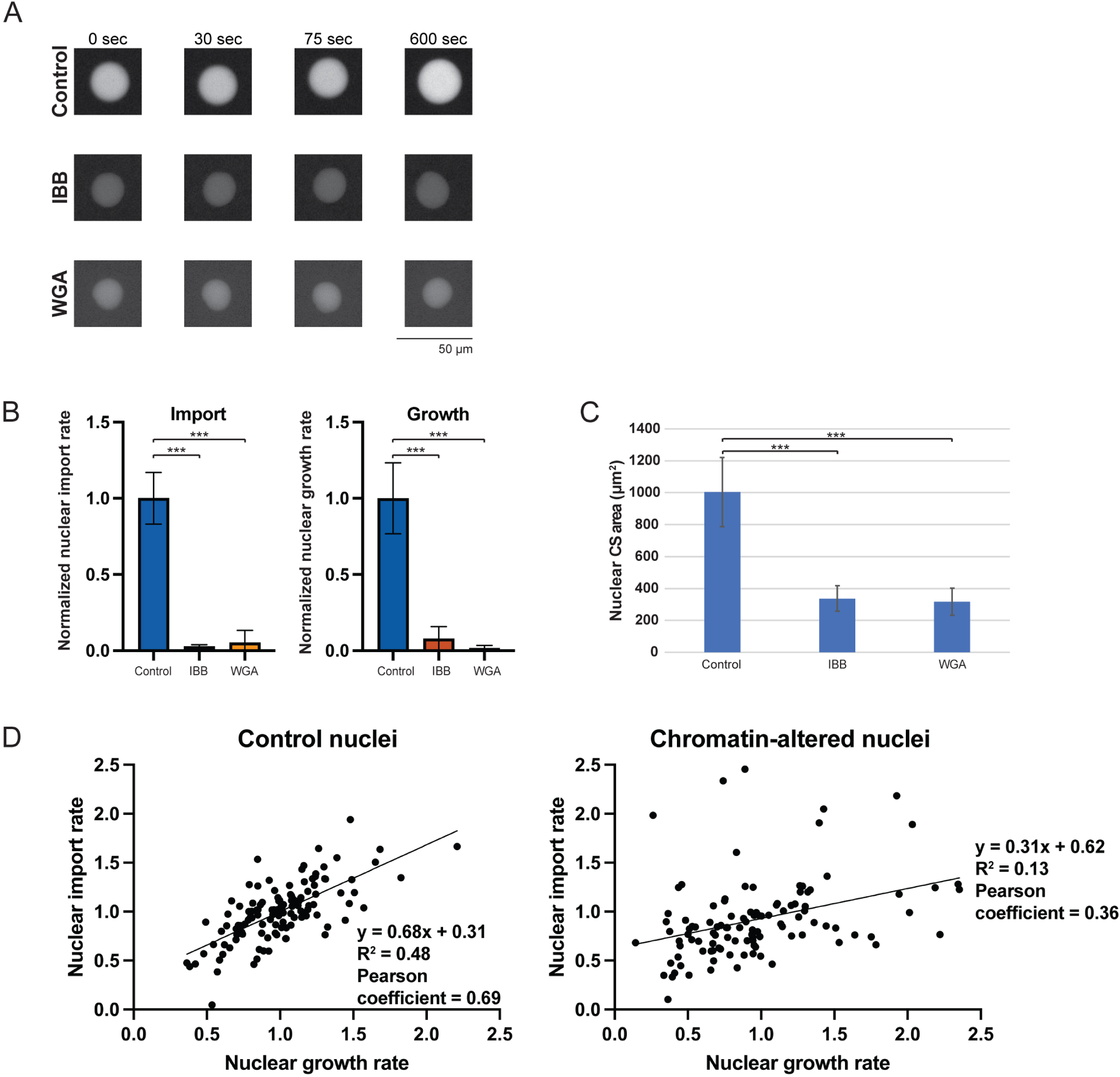
Importin α/μ–mediated nuclear import is necessary for nuclear growth in *X. laevis* egg extract. **(A-B)** Nuclei were assembled in *X. laevis* egg extract with *X. laevis* sperm chromatin. Extract was then supplemented with 28 µM IBB, 0.2 mg/ml WGA, or an equivalent volume of buffer (Control) and incubated for 10 minutes. Then, 2.9 µM GFP-NLS was added and incubated for 5 minutes. Live images were acquired at 15-sec intervals with the same exposure time. **(A)** Time points after GFP-NLS addition are shown for representative time-lapses. Scale bar, 50 µm. **(B)** The initial nuclear import rate was quantified over 1.5 minutes as described in the Methods and normalized to controls. Nuclear growth rate was quantified over 10 minutes as described in the Methods and normalized to controls. At least 8 nuclei were quantified per condition. **(C)** Nuclei were assembled in *X. laevis* egg extract with *X. laevis* sperm chromatin. Extract was then supplemented with 28 µM IBB, 0.2 mg/ml WGA, or an equivalent volume of buffer (Control). After a 1-hour incubation, the nuclei were fixed and visualized by immunofluorescence using mAb414 against the NPC. Nuclear cross-sectional (CS) areas were then determined based on NPC staining. At least 78 nuclei were quantified per condition. **(D)** Nuclear import rate versus nuclear growth rate was plotted for individual nuclei from multiple experiments. Compiled data for control nuclei are from Figs. 2B, 3B, 3D, 3F, 3H, and S1B (n = 123). Compiled data for chromatin-altered nuclei with both increased and decreased nuclear size are for nuclei treated with MNase, VPA, DZNep, Methylstat, and NDMA from Figs. 2B, 3B, 3D, 3F, and 3H (n = 109). Comparing the slopes of the two linear regression lines in GraphPad Prism indicated that the differences between the slopes are extremely significant (p = 0.0008, two-tailed analysis of covariance). One-way ANOVA tests: ***p < 0.001. Error bars represent SD.

**Figure S2.**
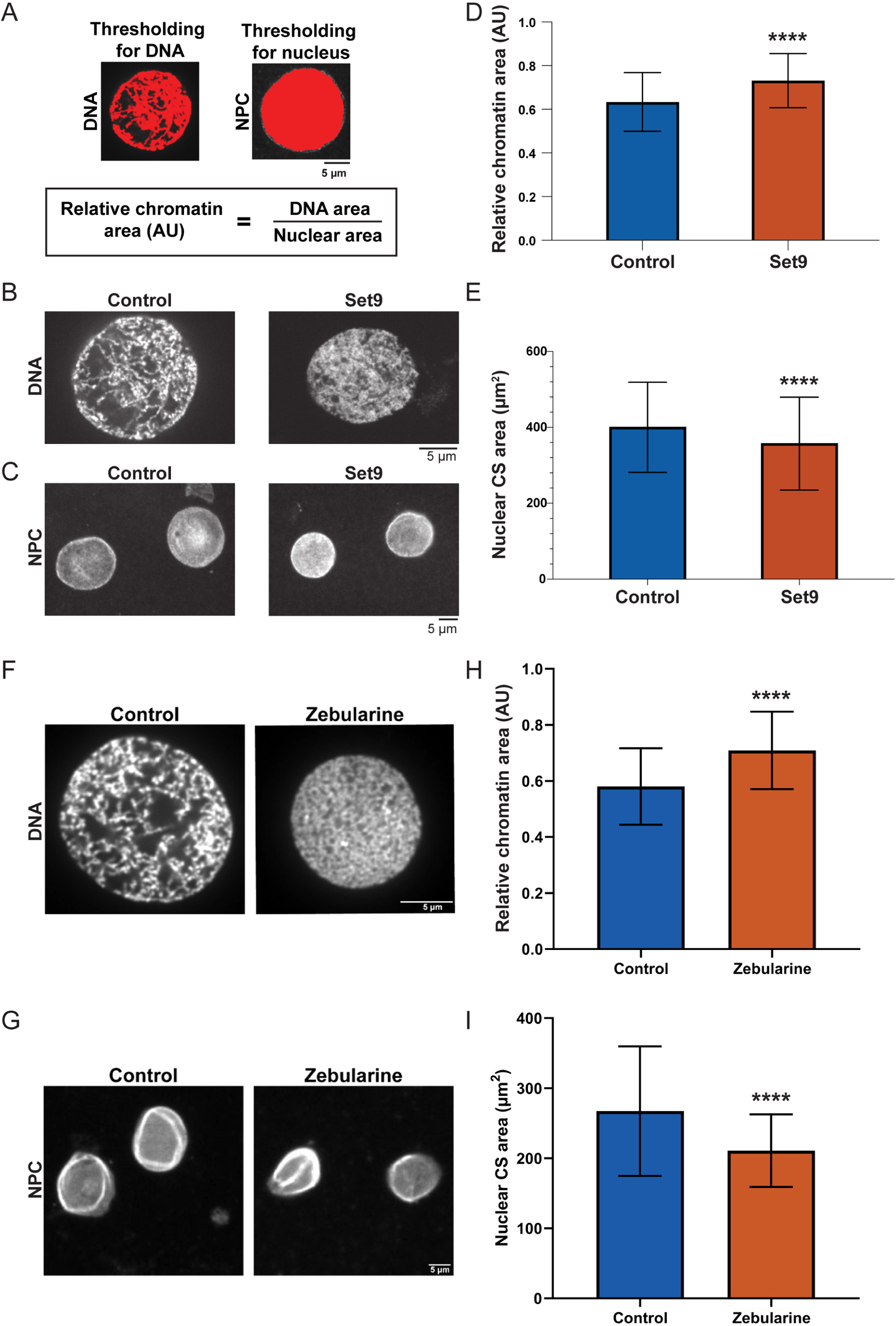
Nuclear size is sensitive to histone and DNA methylation. *X. laevis* egg extracts were supplemented with 1 µM Set9, 100 µM Zebularine, or an equivalent volume of appropriate buffer (Control). *X. laevis* sperm chromatin was then added to induce nuclear assembly. Nuclei were fixed and stained for the NPC (mAb414) or DNA (Hoechst or EthD-2). **(A)** Thresholding was applied to a representative nucleus stained for DNA or NPCs (red). The thresholded DNA area was divided by the thresholded total nuclear CS area to calculate a parameter we refer to as the “relative chromatin area.” Higher values represent less-compact chromatin, whereas lower values represent more-compact chromatin. Scale bar, 5 µm. **(B)** Representative images of Hoechst-stained nuclei are shown. Scale bar, 5 µm. **(C)** Representative images of NPC-stained nuclei are shown. Scale bar, 5 µm. **(D)** Relative chromatin area was quantified for 73 control nuclei and 59 Set9 nuclei from three independent experiments. **(E)** Nuclear CS area was quantified based on NPC staining for 919 control nuclei and 1177 Set9 nuclei from five independent experiments. **(F)** Representative images of EthD-2–stained nuclei are shown. Scale bar, 5 µm. **(G)** Representative images of NPC-stained nuclei are shown. Scale bar, 5 µm. **(H)** Relative chromatin area was quantified for 81 control nuclei and 58 Zebularine-treated nuclei from two independent experiments. **(I)** Nuclear CS area was quantified for 260 control nuclei and 235 Zebularine-treated nuclei from three independent experiments. Two-tailed Student’s t-tests assuming equal variances: ****p < 0.0001. Error bars represent SD.

## MOVIE LEGENDS

**Movie 1. Nuclear growth in *X. laevis* egg extract with or without Benzonase.** Nuclei were assembled in *X. laevis* egg extract with *X. laevis* sperm. Nuclei were left untreated (left panel) or treated with 2.5 U/µL Benzonase (right panel) for 10 minutes, then 2.9 µM GFP-NLS was added. After an additional 5-minute incubation, images were acquired live at 2-minute intervals with the same exposure time. The movie spans ∼120 minutes. Scale bar, 50 μm.

**Movie 2. Nuclear growth in *X. laevis* egg extract with or without micrococcal nuclease (MNase).** Nuclei were assembled in *X. laevis* egg extract with *X. laevis* sperm. Nuclei were left untreated (left panel) or treated with 40 U/µL MNase (right panel) for 10 minutes, then 2.9 µM GFP-NLS was added. After an additional 5-minute incubation, images were acquired live at 2-minute intervals with the same exposure time. The movie spans ∼140 minutes. Scale bar, 50 μm.

**Movies 3-4. Time-dependent compaction of chromatin near the NE.** Nuclei were assembled in *X. laevis* egg extract with *X. laevis* sperm. Nuclei were incubated with 1 µg/ml Hoechst for 5 minutes, and then images were acquired live at 15-sec intervals with the same exposure time. Movie 3 spans 11 minutes and Movie 4 spans 15 minutes. Scale bar, 10 μm.

**Movies 5-8. Chromatin and Lamin B3 dynamics in growing nuclei.** Nuclei were assembled in *X. laevis* egg extract with *X. laevis* sperm. Hoechst (1 µg/mL) and GFP-Lamin B3 (81.6 nM) were added. Following a 5-minute incubation, images were acquired live at 15-sec intervals with the same exposure time. From left to right: GFP-Lamin B3, Hoechst, and merged images (Lamin B3: green and Hoechst: blue). Each movie spans ∼10 minutes. Scale bar, 10 μm.

**Movie 9. The chromatin and lamina concomitantly collapse in rupturing nuclei.** Nuclei were assembled in *X. laevis* egg extract with *X. laevis* sperm. Hoechst (1 µg/mL) and GFP-Lamin B3 (44.5 nM) were added. Following a 5-minute incubation, images were acquired live at 15-sec intervals with the same exposure time. From left to right: GFP-Lamin B3, Hoechst, and merged images (Lamin B3: green and Hoechst: blue).

The movie spans 10 minutes. Scale bar, 10 μm.

